# Design and specificity of long ssDNA donors for CRISPR-based knock-in

**DOI:** 10.1101/178905

**Authors:** Han Li, Kyle A. Beckman, Veronica Pessino, Bo Huang, Jonathan S. Weissman, Manuel D. Leonetti

## Abstract

CRISPR/Cas technologies have transformed our ability to manipulate genomes for research and gene-based therapy. In particular, homology-directed repair after genomic cleavage allows for precise modification of genes using exogenous donor sequences as templates. While both single-stranded DNA (ssDNA) and double-stranded DNA (dsDNA) forms of donors have been used as repair templates, a systematic comparison of the performance and specificity of repair using ssDNA versus dsDNA donors is still lacking. Here, we describe an optimized method for the synthesis of long ssDNA templates and demonstrate that ssDNA donors can drive efficient integration of gene-sized reporters in human cell lines. We next define a set of rules to maximize the efficiency of ssDNA-mediated knock-in by optimizing donor design. Finally, by comparing ssDNA donors with equivalent dsDNA sequences (PCR products or plasmids), we demonstrate that ssDNA templates have a unique advantage in terms of repair specificity while dsDNA donors can lead to a high rate of off-target integration. Our results provide a framework for designing high-fidelity CRISPR-based knock-in experiments, in both research and therapeutic settings.

**Update: November 12th, 2019**
Dear bioRxiv community,
The conclusions of this pre-print (originally posted in August 2017) are outdated. While the experiments we present here are accurate, a recent and more systematic analysis revealed that the integration outcomes driven by different forms of HDR donors are more complex than our methods could originally identify. We initially analyzed donor integration only in FACS-selected cells, which under-estimates alleles where the mis-integration of payload leads to non-functional selection markers, and we quantified integration by ddPCR, which is an indirect read-out of sequence properties. These approaches could not capture the full details of donor integration events in our experiments. To address this, we have now developed a new framework based on long-read amplicon sequencing and an integrated computational pipeline to precisely analyze knock-in repair outcomes across a wide range of experimental parameters. Our new data uncover a complex repair landscape in which both single-stranded and double-stranded donors can lead to high rates of imprecise integration in some cell types. Please read our new bioRxiv pre-print entitled “Deep profiling reveals substantial heterogeneity of integration outcomes in CRISPR knock-in experiments” for further information.
I hope that this example highlights one of the powers of pre-prints: the ability to update scientific discussions (and set records straight) as new results are obtained, often fueled by the availability of new technologies.
Please do not hesitate to contact me directly for any questions or comments.

- Manuel Leonetti

## Introduction

Recent developments in gene editing technologies have transformed our ability to manipulate genomes. Programmable site-specific nucleases, in particular CRISPR/Cas systems, introduce double-strand breaks at target genomic locations that can then be engineered by co-opting endogenous DNA repair mechanisms (1). Notably, homology-directed repair (HDR) can use exogenous donor DNA sequences containing homology to the cleaved genomic target as templates to integrate (knock-in) new genetic information in a locus of interest (2). Gene knock-in strategies have wide applications ranging from correcting disease-causing mutations in a clinical context (3, 4) to introducing fluorescent reporters for the study of protein function in a native cell biology setting (5, 6).

Both single-stranded DNA (ssDNA) and double-stranded DNA (dsDNA) forms of donors can act as efficient HDR templates, but the properties of different donor types have not been systematically compared. Rather, the choice of donor type is often dictated by the size of modifications to be introduced (7, 8). ssDNA donors have been mostly used for applications requiring small edits (5, 9–12), for which commercial oligonucleotides ≤ 200 nt are widely available. In contrast, difficulty in generating long ssDNA has curbed a systematic assessment of ssDNA donors for knock-in applications.

Here, we systematically examine the use of ssDNA donors for CRISPR-based knock-in in human cell lines, and compare ssDNA performance to other forms of templates. We first present a simple and robust method for the preparation of long (~2 kb) ssDNA sequences through optimized reverse-transcription of an RNA intermediate. We next show that long ssDNA sequences are highly efficient HDR templates for the integration of gene-sized reporters. Finally, we demonstrate that ssDNA donors have a unique advantage for specificity of integration in a direct comparison with equivalent dsDNA sequences (plasmid or PCR products). In particular, dsDNA donors can be incorporated at high rates at off-target genomic locations, potentially limiting their use for precise genome editing.

## Results

### An improved reverse transcription method for the generation of long ssDNA

To overcome the size-limit of ssDNA generation, we developed an optimized method for the synthesis of long ssDNA. Reverse-transcription (RT) of an RNA intermediate followed by specific hydrolysis of the RNA strand (Figure 1A) is an efficient and scalable method for ssDNA synthesis (5, 13, 14). However, RT of long sequences is challenging and often results in the accumulation of truncated products because most reverse-transcriptases are poorly processive and unable to transcribe past highly structured RNA regions (15). We reasoned that group II intron RT enzymes, which have evolved to process long and highly structured substrates with high fidelity (15), would permit the synthesis of long ssDNA. Indeed, we found that commercial TGIRT-III (derived from the thermophile *Geobacillus stearothermophilus*) allows for the reliable generation of a 2067 nt-long ssDNA template, while commonly used viral RT enzymes performed poorly (Figure 1B). By Sanger sequencing of TGIRT-III ssDNA products, we measured that the combined error rate of the in vitro transcription and RT steps (Figure 1A) leads to 1 mutation introduced every ~3300 nt in the final ssDNA (data not shown).

**Figure 1:**
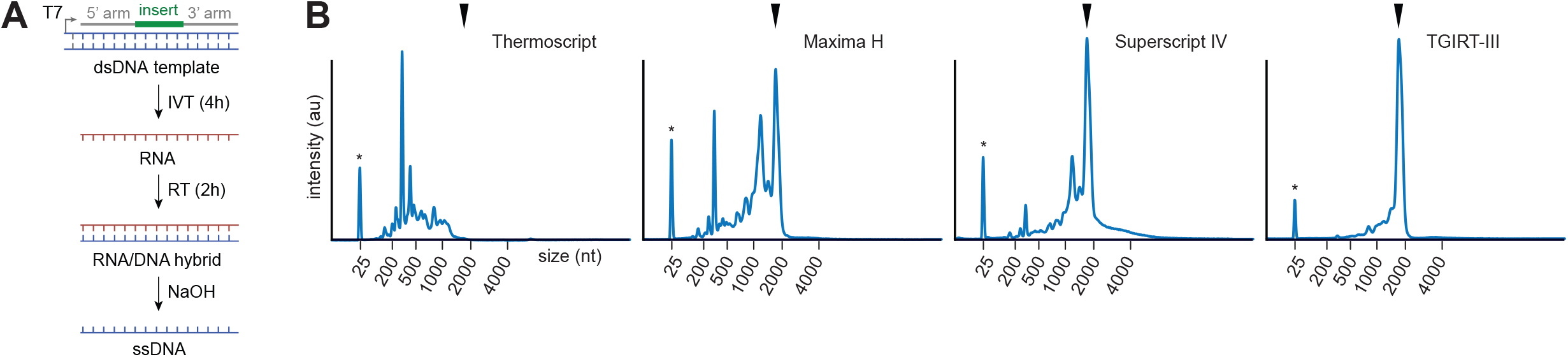
RT-mediated ssDNA synthesis. **(A)** Generation of ssDNA through reverse-transcription (RT) of an RNA intermediate. The RNA strand of the resulting RNA:DNA hybrid can be selectively hydrolyzed at high pH. RNA is first obtained by T7 in vitro transcription (IVT) of a dsDNA template. **(B)** Comparison of RT enzymes for the generation of a 2067 nt-long ssDNA donor. Shown are capillary electrophoresis size profiles of ssDNA measured by Bioanalyzer. Asterisk: 25 nt Bioanalyzer marker. Arrowhead: expected full-size product. TGIRT-III enzyme leads the generation of high-quality ssDNA essentially devoid of truncation products.

Our RT-based ssDNA synthesis scheme can generate >50 μg of final ssDNA (the in vitro transcription step alone resulting in a >100-fold enrichment of material). We routinely carry out all steps in multi-well format and use magnetic beads for nucleic acid purification (16, 17). Initial dsDNA templates (Figure 1A) can be obtained from PCR of sequence-verified plasmids, but we also designed a cloning-free method for the rapid generation of HDR donors (Suppl. Figure 1A). In this method, the three components required to generate the HDR template sequence (5’ and 3’ homology arms and insert) are assembled by fusion PCR of dsDNA fragments. Because non-clonal dsDNA fragments are routinely available by commercial gene synthesis, this enables the cost- and time-efficient generation of HDR templates. Importantly, the same 5’ and 3’ fragments can be used for generating libraries of HDR templates for the integration of different reporter sequences in the same genomic locus (Suppl. Figure 1B).

### Design rules for ssDNA HDR donors

To evaluate the integration efficiency of long ssDNA donor templates, we monitored the knock-in of N- or C-terminal GFP fusion reporters (Figure 2A) using electroporation to deliver *S. pyogenes* Cas9/sgRNA ribonucleoproteins (RNP) and HDR donors into human culture cell lines (10, 18). A summary of all experimental conditions is presented in Table S1. Donors containing ~400-600 nt homology arms lead to ~20-40% GFP knock-in in the RAB11A, CLTA and HIST2H2BE loci in HEK293T cells (Figure 2B). GFP fluorescence matched the expected localization of the targeted proteins, indicating on-target integration (Figure 2B, bottom panels). To illustrate another application of ssDNA-mediated fluorescent tagging for the study of protein function, we also introduced photo-activatable mEos3.2 (19) into CLTA (clathrin light-chain A) and used STORM super-resolution microscopy to image clathrin-coated pits in endogenously-tagged cells (Figure 2C).

**Figure 2:**
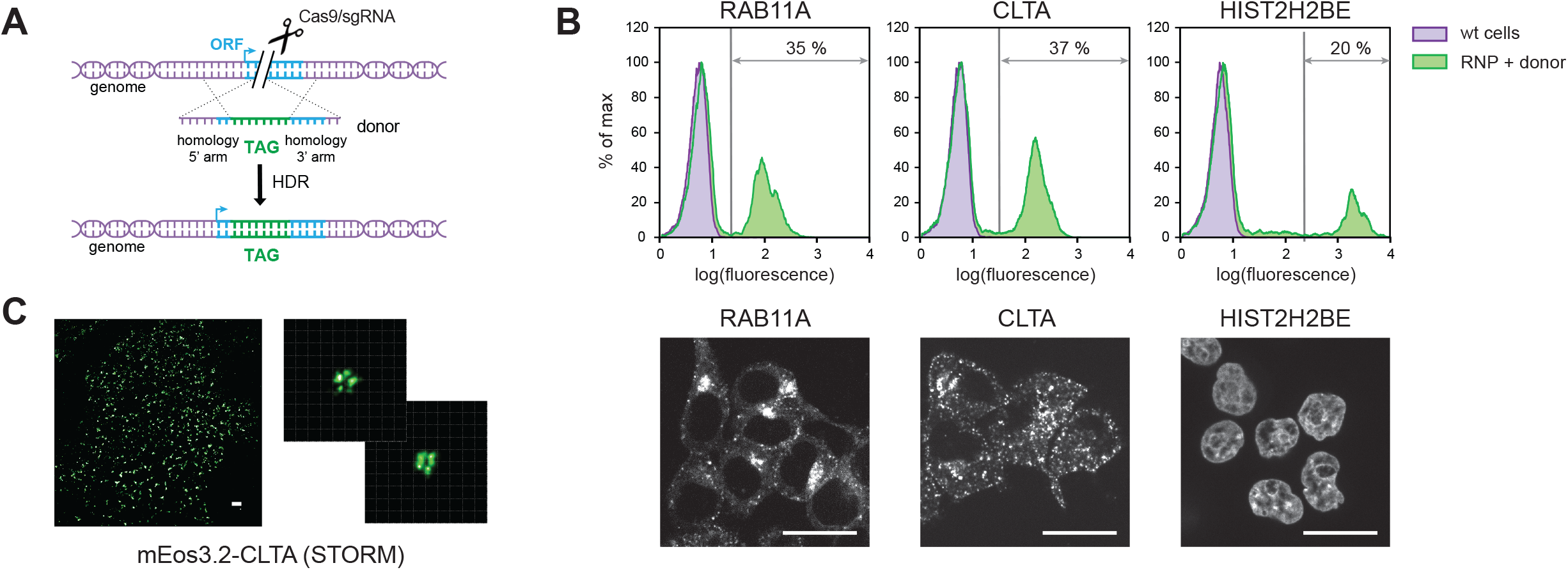
Endogenous gene tagging with ssDNA donors. **(A)** Functional tagging at endogenous genomic loci. Tag (e.g. GFP) is introduced in an endogenous open-reading-frame (ORF) and the resulting fusion protein expressed from the endogenous ORF promoter. **(B)** Endogenous GFP tagging of RAB11A (endosomal Rab protein), CLTA (clathrin light chain) and HISTH2BE (histone) in HEK293T using long ssDNA donors. Knock-in efficiency was measured by flow cytometry analysis ~7 days after Cas9/sgRNA and donor electroporation. Confocal microscopy of GFP-sorted cells is shown (scale bars: 10 μm). **(C)** STORM super-resolution imaging of mEos3.2-CLTA in HEK293T cells. Scale bar: 1 μm; grid size in insert: 100 nm.

We sought to establish rules for maximizing the efficiency of knock-in using long ssDNA donors by systematically characterizing the effects of homology length, amount of donor, donor strand orientation and distance between cutting and insertion sites in our assays. We first prepared donors containing increasingly long homology arms to insert GFP (~700 nt) into RAB11A in HEK293T cells and observed a near-exponential relationship between homology length and knock-in efficiency: longer homology arms drive higher efficiency, with 95% of maximal efficiency reached using ~400 nt arms (Figure 3A). To test whether this rule was specific to a particular locus or a particular insert, we targeted the SEC61B locus with a smaller GFP11 fragment (~60 nt), which we previously leveraged for the high-throughput GFP tagging of proteins in HEK293T cells (5). Interestingly, the relationship between homology length and knock-in efficiency was almost identical to full-length GFP integration in RAB11A (95% maximal efficiency reached using ~300 nt arms, Figure 3B). To verify this result in another human cell line, we repeated the RAB11A GFP integration in K562 cells. GFP was integrated in K562 at much lower rates than in HEK293T but the relationship between increased homology length and efficiency was conserved overall (95% maximal efficiency reached using ~700 nt arms, Figure 3C). From these results, we conclude that ssDNA templates containing long (~400-700 nt) homology arms are optimal donors.

**Figure 3:**
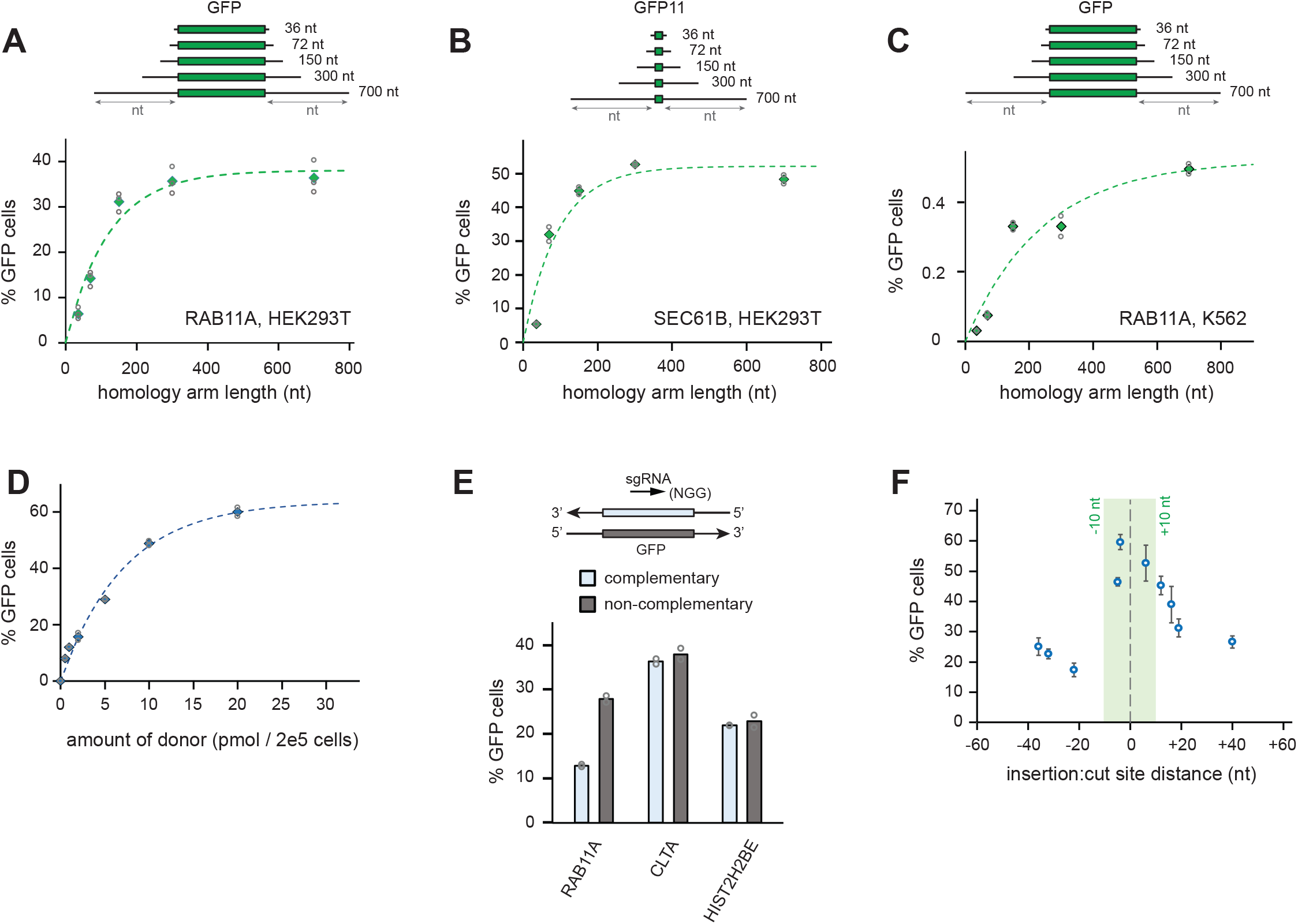
Optimization of ssDNA donor design. **(A)** GFP integration into RAB11A in HEK293T cells using donors of increased homology length (symmetrical donors, length of each arms is shown). Knock-in efficiency is measured by % of GFP-positive cells. Solid markers: average of n=3 independent experiments (individual measurements shown as open circles). An exponential fit is shown (exponential constant = 120 nt). **(B)** GFP11 integration into SEC61B in HEK293T cells. Solid markers: average of n=2 independent experiments (individual measurements shown as open circles). An exponential fit is shown (exponential constant = 90 nt). **(C)** GFP integration into RAB11A in K562 cells. Solid markers: average of n=2 independent experiments (individual measurements shown as open circles). An exponential fit is shown (exponential constant = 240 nt). **(D)** GFP integration into RAB11A in HEK293T cells using increasing amounts of a 300 nt homology arms donor. 100 pmol Cas9 RNP and 2×10^5^ cells were used for all samples. Solid markers: average of n = 2 independent experiments (individual measurements shown as open circles). **(E)** Effect of ssDNA donor strand orientation. GFP integration into RAB11A, CLTA and HIST2H2BE into HEK293T cells. Average of n=2 independent experiments (individual measurements shown as open circles. Respective orientation of sgRNA and donor strand is shown. **(F)** Effect of distance between integration site and site of sgRNA cleavage. 2×GFP11 integration into SEC61B in HEK293T cells. Error bars represent s.e.m of n = 4-5 independent experiments.

We next characterized the amount of donor needed for maximal knock-in rate. In HEK293T cells, 95% maximal efficiency was obtained using ~20 pmol of donor for 2×10^5^ cells (Figure 3D). We also tested whether the strand orientation of ssDNA donors impacted knock-in efficiency. Comparing donors complimentary or non-complimentary to sgRNAs targeting RAB11A, CLTA and HISTH2BE in HEK293T cells, we observed no consistent difference between ssDNA orientation and knock-in efficiency, although the sgRNA-complementary orientation was a poorer template for RAB11A (Figure 3E). Finally, we characterized how the rate of reporter integration varied with its distance to the site of double-strand cleavage. We designed 10 sgRNAs leading to cleavage between −36 and +40 nt of a single SEC61B insertion site in HEK293T cells. Following the integration of a 2×GFP11 sequence (165 nt), we observed maximal efficiency using sgRNAs cutting within ±10 nt of the integration site (Figure 3F), confirming other reports (20, 21).

### Systematic comparison of different forms of HDR donors for efficiency and specificity

Finally, we systematically compared the performance of ssDNA, plasmid (non-linearized) or PCR-derived donors for GFP knock-in in human cell lines. Using equimolar GFP donors (150 nt arms) targeting RAB11A and CLTA in HEK293T cells, we observed that PCR donors performed best in terms of apparent efficiency (as estimated by % GFP-positive cells), followed by ssDNA and plasmid donors (Figure 4A-B). Next, we characterized the specificity of donor integration. To measure knock-in specificity at the intended target site, we designed a digital droplet PCR (ddPCR) assay to compare the total frequency of GFP integration in the genome with the frequency of GFP integration at the site of interest (see design in Figure 4C). We defined an “on-target” integration percentage by the ratio of [target-specific GFP integration]:[total GFP integration]. We first benchmarked this assay using genomic DNA (gDNA) from control wild-type HEK293T cells spiked with different ratios of plasmids containing “on-target” or “off-target” sequence contexts (Figure 4D). We observed a 1:1 linear relationship between plasmid ratios and on-target percentage measured by ddPCR, validating our assay.

**Figure 4:**
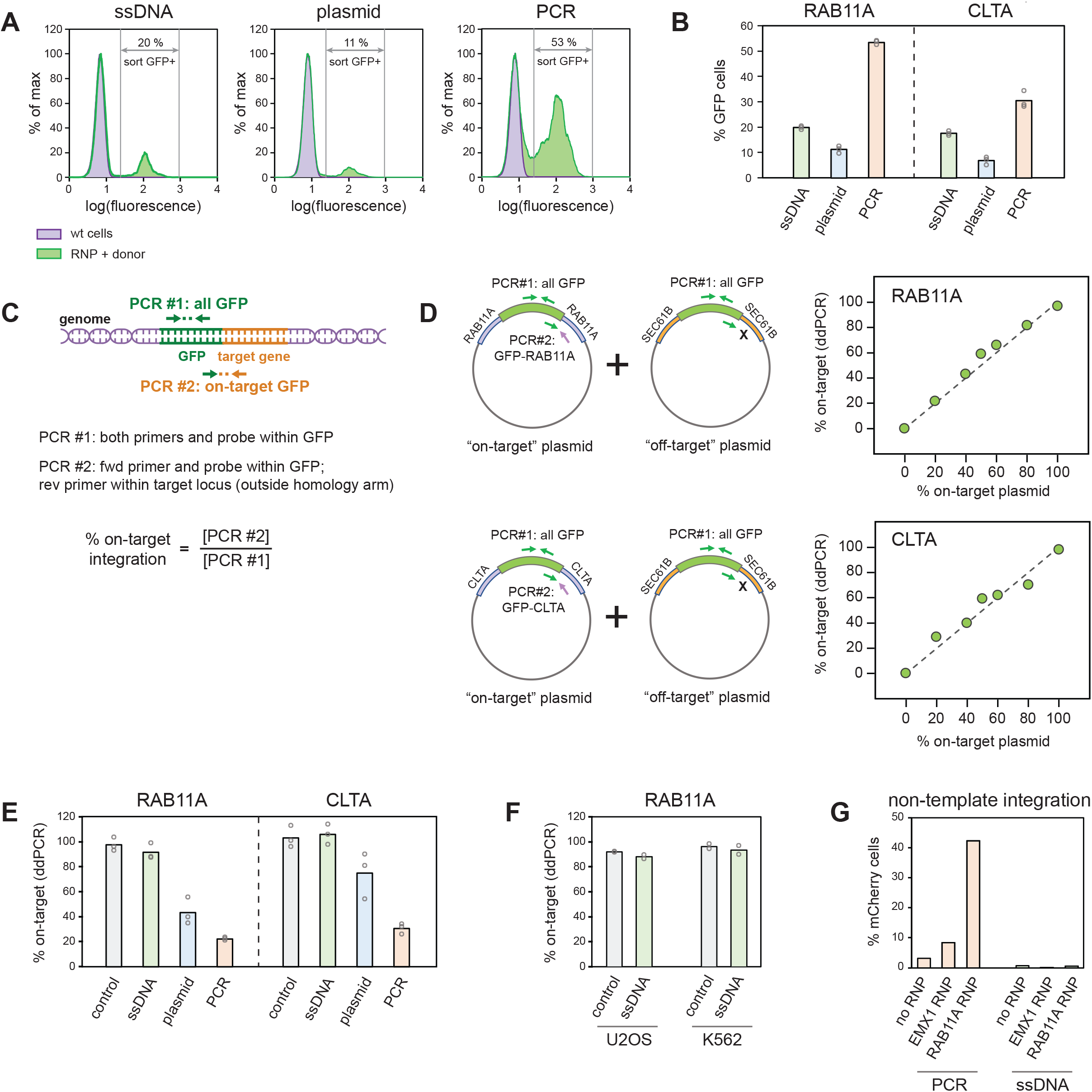
Comparison between ssDNA, plasmid and PCR donors. **(A)** Efficiency of GFP integration into RAB11A in HEK293T cells as measured by flow cytometry. **(B)** GFP knock-in efficiencies into RAB11A and CLTA in HEK293T cells. Average of n = 3 independent experiments (individual measurements shown as open circles). **(C)** Principle of ddPCR-based measurement of on-target GFP integration. **(D)** Validation of ddPCR on-target measurement using control plasmids spiked into wild-type HEK293T genomic DNA. Each set of experiments contains an “on-target” plasmid (GFP into the targeted locus, RAB11A or CLTA) and an “off-target” plasmid (GFP into the non-targeted SEC61B locus), mixed in different ratios. On-target ddPCR measurement as defined in is plotted against the ratio of on-target to off-target plasmid. Dotted line represents the y = x diagonal. **(E)** Measurement of on-target integration of GFP into RAB11A and CLTA in HEK293T cells (same samples as in (B), analysis performed on GFP-positive sorted populations). Average of n = 3 independent experiments (individual measurements shown as open circles). Positive controls: wild-type gDNA spiked with “on-target” control plasmids as in (D). **(F)** Measurement of on-target integration of GFP into RAB11A in U2OS and K562 cells (analysis performed on GFP-positive sorted populations). Average of n = 2 independent experiments (individual measurements shown as open circles). Positive controls: wild-type gDNA spiked with “on-target” control plasmids as in (D). **(G)** Non-template integration of CMV-mCherry devoid of homology arms in HEK293T cells. CMV-mCherry donors were electroporated without Cas9 or with RNP targeting EMX1 or RAB11A.

To measure integration specificity, we sorted GFP-positive cells and prepared gDNA ~30 days after RNP/donor electroporation (to allow for clearance of non-integrated donor molecules that could otherwise interfere with our measurements). We included gDNA from wild-type cells spiked with “on-target” plasmids as positive controls. Surprisingly, different forms of HDR donors lead to vastly different levels of integration specificity (Figure 4E). While ssDNA donors exhibited specificity levels comparable to positive controls, dsDNA donors showed high levels of off-target integration for both RAB11A and CLTA targeting. PCR donors performed much worse than plasmid, showing only ~20-30% specific integration (Figure 4E). Our GFP fusion inserts do not include a promoter, so that off-target integrants would most likely not drive GFP expression. Finally, to verify the specificity of ssDNA-mediated integration in other human cell lines, we repeated RAB11A GFP targeting in U2OS and K562 cells and confirmed on-target levels comparable to positive controls (Figure 4F).

We hypothesized that the high rate of non-specific dsDNA knock-in originates from non-homologous integration of the donors at unwanted sites of double-strand breaks, which would be avoided with ssDNA. To test this, we followed the non-template integration of a CMV-mCherry construct that drives mCherry expression regardless of integration site. This construct did not contain any homology to the human genome and therefore could not drive HDR-based knock-in. Delivering CMV-mCherry in HEK293T cells as either ssDNA or PCR product, in the presence or absence of Cas9 RNPs targeting arbitrarily chosen loci, we observed significant non-template insertion of PCR, but not ssDNA donors (Figure 4G; mCherry fluorescence was measured 28 days post-electroporation to allow for clearance of non-integrated donor molecules). These results suggest that, as opposed to dsDNA, ssDNA is not a substrate for non-homologous integration and is therefore a more specific knock-in template.

## Discussion

Altogether, our results support three main conclusions. First, we report an optimized method for the generation of long ssDNA templates, paving the way for the use and analysis of ssDNA for large knock-in insertions. Direct strand-specific digestion of dsDNA is an attractive alternative to our RT method (13), but in our experience generates only < 10-fold lower amounts of material. Importantly, our data shows that efficient knock-in by electroporation requires high pmol quantities of donor (Figure 3D) and therefore necessitates large amounts of material. The amounts required for other applications, for example direct injection into mouse zygotes, are much lower (8, 14) and could be well suited to alternative methods for ssDNA synthesis. Single-stranded recombinant adeno-associated virus constructs up to ~4.5 kb in length can also be used but require fairly involved preparation protocols (22, 23).

Second, our results delineate general guidelines for the design of ssDNA donors. We propose that ssDNA donors containing ~500 nt homology arms, non-complementary to the sgRNA used, and preferably coupled with a sgRNA cutting within 10 nt of the desired integration site are optimal knock-in templates. In particular, we find that long homology is advantageous across all of the loci, insert sizes and cell lines we tested. Similar homology requirements have also been described in studies using dsDNA HDR templates (24, 25). Homology length and amount of donor most likely act in concert to drive knock-in efficiency. We routinely use ssDNA oligonucleotide donors containing short 50-70 nt homology arms for the integration of GFP11 fragments in cells and observe very high knock-in efficiencies (5). In our experience, short oligonucleotides can be delivered in much higher molar quantities than long ssDNA donors before toxicity arises, and can therefore drive efficient knock-in despite shorter homology. Of note, we found variable ssDNA-mediated integration efficiencies in HEK293T and K562 cells (compare Figures 3A and 3C). The HDR pathway used by ssDNA templates is still unclear but likely differs from classical RAD51-dependent repair (21, 26–28), and might be active at different levels across cell lines.

Finally, our results demonstrate a unique advantage of ssDNA donors in limiting off-target integration events. While non-template integration of donors in gene editing experiments is a known problem (22, 29), few studies have directly quantified the occurrence of off-target integration events (6, 30). A recent study in humans stem cells reported prevalent random integration of plasmid donors and emphasized the need for careful characterization of knock-in cells (6). Our data show that plasmid or PCR donors can lead to high rates of off-target integration, which might limit their use for precise genome engineering. In contrast, our results in HEK293T cells demonstrate that ssDNA donors can drive integration of large inserts in a manner that is both highly efficient and very specific. We also show that the specificity of ssDNA-mediated knock-in is preserved amongst multiple cell lines. Based on these findings, we believe that ssDNA donors are advantageous for high-precision genome engineering applications. The synthesis and design strategies we have highlighted should encourage their broader adoption, in both research and therapeutic settings.

## Material and methods

### Cell culture

HEK293T and U2OS cells were cultured in high-glucose DMEM supplemented with 10% FBS, 1 mM glutamine and 100 μg/mL penicillin/streptomycin (Gibco). K562 cells were cultured in RPMI 1640 medium containing 25 mM Hepes and supplemented with 10% FBS, 1 mM glutamine and 100 μg/mL penicillin/streptomycin (Gibco). HEK293T cells constitutively expressing a GFP1-10 construct were prepared as in (5). All cell lines were mycoplasma-free and routinely tested.

### Nucleic acid purification with magnetic beads

Magnetic beads for nucleic acid purification were prepared as in (17). For a 50 mL working solution of beads, 1 mL carboxylate-modified magnetic bead solution (GE Healthcare #65152105050250) is first washed with 3×1 mL RNAse-free TE buffer (10 mM Tris, 1 mM EDTA, pH 8.0) on a magnetic stand. The bead pellet is then resuspended in DNA precipitation buffer (RNAse-free): {1 M NaCl; 10 mM Tris-HCl pH 8.0; 1 mM EDTA; 18% w/v PEG8000 (Sigma BioUltra); 0.05% v/v Tween20 (Sigma)}. For nucleic acid precipitation and purification, magnetic beads in precipitation solution are added to nucleic acid samples in appropriate ratios and incubated for 10 min at RT. Bead-bound nucleic acids are then isolated on a magnet and washed 2× in RNAse-free 70% EtOH. Beads are air-dried (5-10 min at RT) and the nucleic acids eluted in RNAse-free H_2_O.

### sgRNA in-vitro transcription

sgRNA is obtained by in vitro transcription of a DNA template of the following sequence: {5’- TAA TAC GAC TCA CTA TAG NN NNN NNN NNN NNN NNN NNG TTT AAG AGC TAT GCT GGA AAC AGC ATA GCA AGT TTA AAT AAG GCT AGT CCG TTA TCA ACT TGA AAA AGT GGC ACC GAG TCG GTG CTT TTT TT-3’} containing a T7 promoter (TAATACGACTCACTATA), a gene-specific 19-nt sgRNA protospacer sequence preceded by a G for T7 transcription (GNNNNNNNNNNNNNNNNNNN) and a common sgRNA constant region. The DNA template is generated by overlapping PCR using a set of 4 primers: 3 primers common to all reactions (forward primer ML557: 5’- TAA TAC GAC TCA CTA TAG -3’; reverse primer ML558: 5’- AAA AAA AGC ACC GAC TCG GTG C -3’ and reverse primer ML611: 5’- AAA AAA AGC ACC GAC TCG GTG CCA CTT TTT CAA GTT GAT AAC GGA CTA GCC TTA TTT AAA CTT GCT ATG CTG TTT CCA GCA TAG CTC TTA AAC -3’) and one gene-specific primer (forward primer 5’- TAA TAC GAC TCA CTA TAG NNN NNN NNN NNN NNN NNN NG TTT AAG AGC TAT GCT GGA A -3’). A 40 μL PCR is set using Kapa HotStart HiFi reagents (Kapa Biosystems #KK2601) containing 40 pmol ML557, 40 pmol ML558, 0.8 pmol ML611 and 0.8 pmol gene-specific primer. PCR reaction is amplified in a thermocycler: 95°C for 3 min, 20 cycles of {98°C for 20 s, 57°C for 15 s, 72°C for 5 s}, 72°C for 1 min, 4°C final. For purification, 60 μL 100% EtOH and 100 μL magnetic beads in precipitation solution were added to the reaction. After magnetic purification, PCR products are eluted in 12 μL of RNAse-free {2 mM Tris-HCl pH 8.0}. 4 μL of purified PCR products are used in a 40 μL IVT reaction using HiScribe T7 polymerase (NEB #E2040S) and containing 5 mM each NTPs (NEB # N0466S). After incubation for 3 hours at 37°C, 1 μL TurboDNAse (Thermo #AM2238) is added to the reaction and incubated 15 min at 37°C. For purification, 80 μL 100% EtOH and 100 μL magnetic beads in precipitation solution are added to the reaction. After magnetic purification, sgRNA products are eluted in 20 μL of RNAse-free H_2_O and the concentration adjusted to 130 μM (typical yield: 4000 pmol sgRNA). sgRNA quality is routinely checked by running 3 pg of the purified sgRNA on a 10% polyacrylamide gel containing 7M urea (Novex TBE-Urea gels, Thermo).

### Fusion PCR of HDR templates

Details of fusion PCR design are given in Suppl. Figure 1 and in Table S1 (sheet 2). For fusion PCR, gene fragments were purchased from IDT (gBlocks). PCR primers: ML1125 (5’- GGG AAC CTC TTC TGT AAC TCC TTA GC -3’) and ML1126 (5’- CCT GAG GGC AAA CAA GTG AGC AGG -3’). A 100 μL PCR reaction is set using Kapa HotStart HiFi reagents (Kapa Biosystems #KK2601) containing 125 pmol ML1125, 125 pmol ML1126, 16 fmol 5’ arm fragment, 16 fmol 3’ arm fragment and 80 fmol of insert fragment. PCR reaction is amplified in a thermocycler: 95°C for 3 min, 30 cycles of {98°C for 20 s, 69°C for 15 s, 72°C for 15 s/kb}, 72°C for 5 min, 4°C final. For purification, 60 μL magnetic beads in precipitation solution are added to the reaction. After magnetic purification, PCR products are eluted in 15 μL of RNAse-free {2 mM Tris-HCl pH 8.0}.

### ssDNA generation: step 1 – PCR

All constructs used here are amplified by PCR of sequence-verified plasmids, except for the examples of fusion PCR shown in Suppl. Figure 1. Fusion PCR products can be used identically. A 100 μL PCR reaction is set using Kapa HotStart HiFi reagents (Kapa Biosystems #KK2601) containing 125 pmol ML1125, 125 pmol ML1126 and 10 ng plasmid template. PCR reaction is amplified in a thermocycler: 95°C for 3 min, 30 cycles of {98°C for 20 s, 69°C for 15 s, 72°C for 15 s/kb}, 72°C for 5 min, 4°C final. 2 μL DpnI (NEB # R0176L) is then added and the reaction incubated 30 min at 37°C. For purification, 60 μL magnetic beads in precipitation solution are added to the reaction. After magnetic purification, PCR products are eluted in 15 μL of RNAse-free {2 mM Tris-HCl pH 8.0}.

### ssDNA generation: step 2 – IVT

A 100 μL IVT reaction is set using HiScribe T7 polymerase (NEB #E2040S). The reaction contained: 10 μL 10× T7 buffer, 2 μL {0.1 M DTT}, 50 μL NTP mix (25 mM each, NEB #N0466S), 2 μL RNAse.In (Promega #N2115), 20 μL RNAse-free H_2_O, 6 μL DNA from step 1 and 10 μL T7 HiScribe enzyme. The reaction is incubated 3 hours at 37°C, after which 4 μL TurboDNAse (Thermo #AM2238) is added to the reaction and incubated 15 min at 37°C. For purification, 120 μL magnetic beads in precipitation solution are added to the reaction. After magnetic purification, RNA products are eluted in 120 μL of RNAse-free H_2_O. This step can be extremely efficient (yield can exceed 1 mg RNA). For high-yield reactions, magnetic purification can be challenging because RNA amount exceeds the binding capacity of the beads. In such cases, the reaction can be scaled down or more beads added.

### ssDNA generation: step 3 – RT

The RT reaction is set up as follows. First, 50 μL of RNA (maximum: 500 pmol) is mixed with 8 μL of gene-specific RT primer (100 μM in H_2_O) and 12 μL dNTP mix (25 mM each, Thermo #1122). To allow for primer annealing, the reaction is incubated 5 min at 65°C and placed immediately on ice for another 5 min. Then are added: 20 μL 5x RNAse-free RT buffer {250 mM Tris-HCl pH 8.3, 375 mM KCl, 15 mM MgCl_2_}, 5 μL {0.1 M DTT}, 1 μL RNAse.In (Promega #N2115) and 5 μL TGIRT-III enzyme (InGex). The reaction was incubated 1.5 hours at 58°C, after which RNA is hydrolyzed by the addition of 42 μL {0.5 M NaOH, 0.25 M EDTA pH 8.0} and incubated 10 min at 95°C. NaOH is quenched with 42 μL {0.5 M HCL}. For purification, 280 μL magnetic beads in precipitation solution are added to the reaction. After magnetic purification, ssDNA products are eluted in 25 μL sterile H_2_O. The reaction can be scaled down two-fold to fit in PCR strip format. Typical yields: 50-200 pmol ssDNA per 500 pmol RNA template.

### RNP preparation and electroporation

Cas9/sgRNA RNP complexes were prepared following methods by Lin et al. (10) with some modifications. Cas9 protein (pMJ915 construct, containing two nuclear localization sequences) was expressed in E. coli and purified by the UC Berkeley Macrolab following protocols described by Jinek et al. (31). HEK293T cells were treated with 200 ng/mL nocodazole (Sigma) for 15 hours before electroporation to increase gene editing efficiency as in (10). K562 and U2OS cells were not treated. RNP complexes were assembled with 100 pmol Cas9 protein and 130 pmol sgRNA just prior to electroporation, and combined with HDR template in a final volume of 10 μL. First, 1 μL sgRNA (130 μM stock) is added to 2 μL high-salt RNP buffer {580 mM KCl, 40 mM Tris-HCl pH 7.5, 20% v/v glycerol, 2 mM TCEP-HCl pH 7.5, 2 mM MgCl_2_ RNAse-free} and incubated at 70°C for 5 min. 2.5 μL of Cas9 protein (40 μM stock in Cas9 buffer, ie. 100 pmol) is then added and RNP assembly is carried out at 37°C for 10 min. Finally, HDR templates and sterile RNAse-free H_2_O are added to 10 μL final volume (for experiments in Figure 3B, 5 μg salmon sperm DNA – Thermo #15632011 – was also included in each sample as a neutral carrier to reduce differences in mass of nucleic acid included between samples). Electroporation is carried out in Amaxa 96-well shuttle Nuleofector device (Lonza) using SF- or SE-cell line reagents (Lonza) following the manufacturer’s instructions. Cells are washed with PBS and resuspended to 100 cells/μL in SF solution (HEK293T, K562) or SE solution (U2OS) immediately prior to electroporation. For each sample, 20 μL of cells (ie. 2×105 cells) are added to the 10 μL RNP/template mixture. Cells are immediately electroporated using CM130 (HEK293T, K562) or CM105 (U2OS) programs and transferred to 1 mL pre-warmed culture media in a 24-well plate. Electroporated cells are cultured for > 5 days prior to analysis.

### Flow cytometry and analysis

Analytical flow cytometry was carried out on a LSR II instrument (BD Biosciences) and cell sorting on a FACSAria II (BD Biosciences). Flow cytometry data analysis and figure preparation was done using the FlowJo software (FlowJo LLC).

### Imaging

For confocal microscopy, cells were grown in glass bottom culture dishes with #1.5 high performance cover glass. Live cells were imaged on an inverted Nikon Ti-E microscope, Yokogawa CSU-22 confocal scanner unit, Plan Fluor 10×/0.3 NA objective or Plan Apo VC 60×/1.4 NA oil objective, an Andor EM-CCD camera (iXon DU897) and Micro-Manager software.

For STORM, cells were plated on a poly-lysine-coated imaging chamber and fixed in 4% PFA. mEos3.2 STORM images were acquired with a 405 nm activation laser (OBIS 405, Coherent), and a 561 nm imaging laser (Sapphire 561 LP, Coherent). All lasers were aligned, expanded, and focused at the back focal plane of the 1.4 NA 100× oil immersion objective (UPlanSApo, Olympus). Images were recorded on an Andor EM-CCD camera (iXon DU897E), and processed with previously established home-written software. The OBIS lasers were controlled directly by the computer, while a quad-band (ZT405/488/561/640rpc, Chroma) dichroic mirror and a band-pass filter (ET595/50m Chroma for 561nm excitation) separated the fluorescence emission from the excitation light. Maximum laser power used during STORM measured before the objective was 17 μW for 405 nm (~0.32 W/cm2 at the sample), and 28 mW for 561 nm (~0.52 kW/cm^2^ at the sample). STORM frames were recorded at a rate of 60 Hz, with an EM-CCD camera gain of 30. During image acquisition, the axial drift of the microscope stage was stabilized by a home-built focus stabilization system utilizing the reflection of an infrared laser off the sample. 40,000 frames were collected per super-resolution sample, and Insight3 software was used to reconstruct and analyze them. Cells were imaged in PBS.

### Digital droplet PCR

GFP-positive cells were isolated by FACS and genomic DNA (gDNA) from ~5×10^6^ cells was prepared using PureLink Genomic DNA purification kits (Thermo #K182001) following manufacturer’s instructions. 1 μg gDNA in 50 μL H_2_O was fragmented to 800 bp in a Covaris M220 sonicator. For experiments using plasmid controls, 0.5 pg of plasmid was spiked into gDNA before fragmentation. For each ddPCR reaction, a 10× primer/probe mix was prepared containing: {9 μM forward primer, 9 μM reverse primer, 2.5 μM probe}. Sequences of all primers and probes can be found in Table S1. Probes were purchased from IDT (ZEN™/Iowa Black™ FQ quenched probes). For each reaction, two sets of primer/probes in separate fluorescent channels (FAM or HEX) were used. In the FAM fluorescent channel, a set of primers and probes amplified either total GFP sequences or GFP sequences at the intended targeted locus (Figure 3C, see details in Table S1). In the HEX channel, primers and probes amplified a genomic sequence >8kb away from the Cas9 target site that served as an internal “housekeeping” normalization measurement for the total amount of gDNA in each ddPCR sample (See Table S1). 22 μL ddPCR mixes were set with: 2.2 μL 10× FAM primer/probe mix, 2.2 μL 10× HEX “housekeeping” primer/probe mix, 5 μL fragmented gDNA (100 ng total), 1.6 μL H_2_O and 11 μL 2× ddPCR Supermix (Bio-Rad #1863024). 20 μL of ddPCR mixes were used to generate droplets in a QX200 Droplet Generator (Bio-Rad), following manufacturer’s instructions. Droplets where then transferred to a 96-well PCR plate and submitted to PCR amplification in a C1000 thermocycler (Bio-Rad) as per manufacturer’s instructions: 95°C for 10 min, then 40 cycles of {94°C for 30 s, 60°C for 45 s, 72°C for 1 min}, 98°C for 10 min, 10°C forever. Amplified droplets were immediately read on QX200 Droplet Reader (Bio-Rad) and amplification results (copies of amplified sequences per well) were analyzed using QuantaSoft Pro Software (Bio-Rad). For each sample, amplicon copies in the FAM channel were normalized to amplicon copies in the “housekeeping” HEX channel to account for differences in total amount gDNA detected in each sample.

## Supplementary Figure legend

**Suppl. Figure 1:**
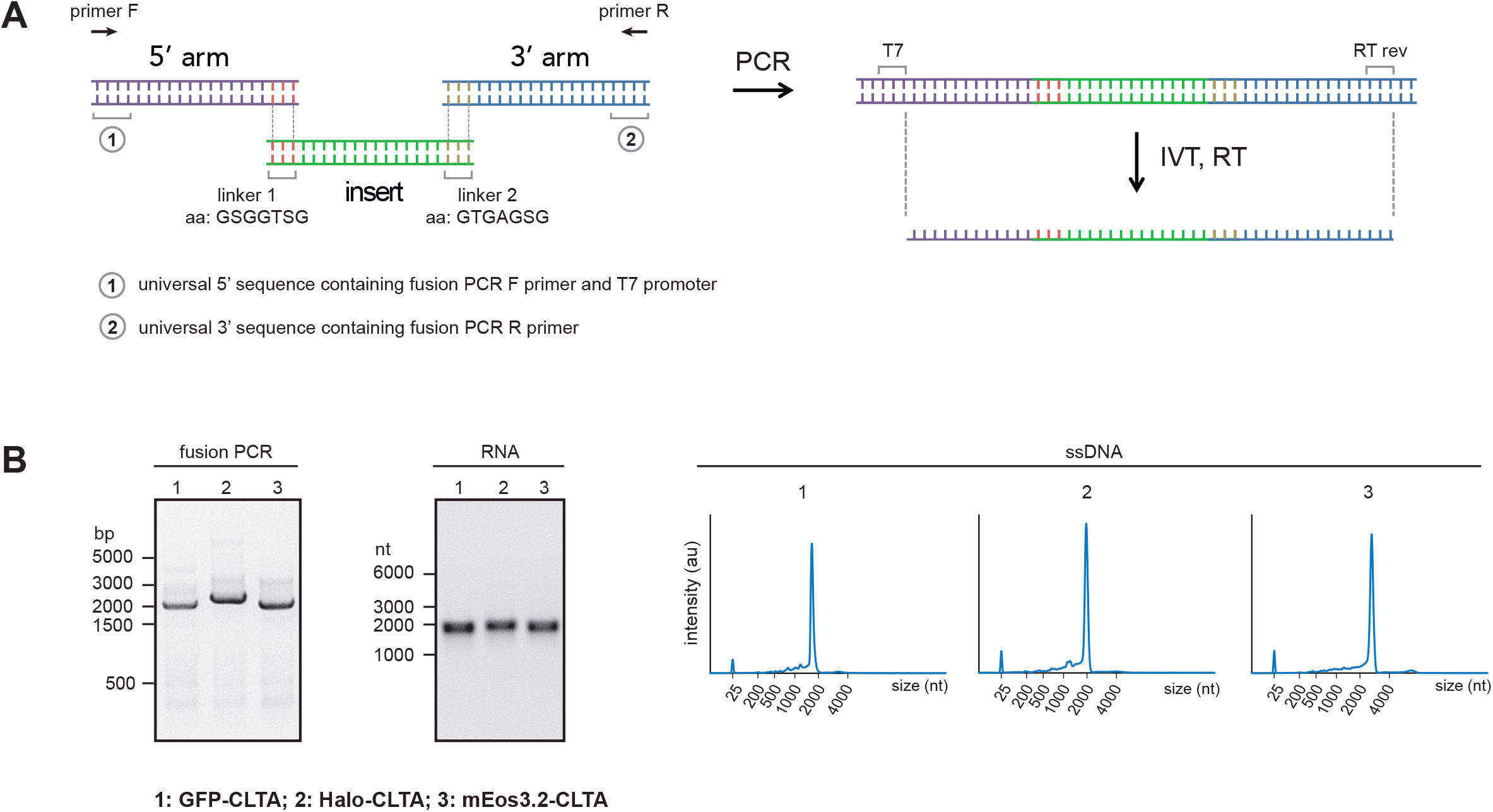
Fusion PCR design for HDR templates. **(A)** Fusion PCR design. Templates are assembled from three independent dsDNA fragments containing 5’ arm, knock-in insert and 3’ arm. Our design is built for the in-frame integration of protein reporters into endogenous genes. Therefore, the overlapping DNA sequences between fusion fragments are designed to translate into amino acid linkers between the target protein and reporter. The amino acid sequence of both linkers is shown. Terminal 5’ and 3’ sequences (1) and (2) contain universal fusion PCR primers binding sites, as well as T7 promoter sequence for IVT. The full-length fusion PCR product can be turned into ssDNA by IVT and RT using the scheme shown in Figure 1A. Gene-specific reverse primers are used for RT (RT rev). **(B)** Example of fusion PCR to prepare templates for the integration of different reporters into CLTA, using the same 5’ and 3’ arm fragments. Three reporters are used: GFP, HaloTag and mEos3.2. For each step in ssDNA generation (Figure 1A), an electrophoresis size profile of the products in shown. Fusion PCR and RNA: agarose gel electrophoresis. ssDNA: Bioanalyzer capillary electrophoresis. RT was carried out using TGIRT-III.

